# Resting state functional connectivity patterns associated with pharmacological treatment resistance in temporal lobe epilepsy

**DOI:** 10.1101/382523

**Authors:** Christina Pressl, Philip Brandner, Stefan Schaffelhofer, Karen Blackmon, Patricia Dugan, Manisha Holmes, Thomas Thesen, Ruben Kuzniecky, Orrin Devinsky, Winrich A. Freiwald

## Abstract

There are no functional imaging based biomarkers for pharmacological treatment response in temporal lobe epilepsy (TLE). In this study, we investigated whether there is an association between resting state functional brain connectivity (RsFC) and seizure control in TLE. We screened a large database containing resting state functional magnetic resonance imaging (Rs-fMRI) data from 286 epilepsy patients. Patient medical records were screened for seizure characterization, EEG reports for lateralization and location of seizure foci to establish uniformity of seizure localization within patient groups. Rs-fMRI data from patients with well-controlled left TLE, patients with treatment-resistant left TLE, and healthy controls were analyzed. Healthy controls and cTLE showed similar functional connectivity patterns, whereas trTLE exhibited a significant bilateral decrease in thalamo-hippocampal functional connectivity. This work is the first to demonstrate differences in neural network connectivity between well-controlled and treatment-resistant TLE. These differences are spatially highly focused and suggest sites for the etiology and possibly treatment of TLE. Altered thalamo-hippocampal RsFC thus is a potential new biomarker for TLE treatment resistance.

**Summary:** Resting State functional magnetic resonance imaging (Rs-fMRI), previously utilized to predict lateralization of seizure foci in temporal lobe epilepsy (TLE), is utilized to determine potential mechanisms and biomarkers for treatment-resistant and well-controlled unilateral TLE. We found significant differences in thalamo-hippocampal functional connectivity between treatment-resistant and well-controlled TLE patients. Differences in functional connectivity were focused to thalamo-hippocampal connections and more pronounced in the hemisphere ipsilateral to seizure foci. Aberrant functional connectivity patterns as measured by Rs-fMRI could thus serve as biomarkers for treatment response in TLE.

## Introduction

Temporal lobe epilepsy (TLE) is the most common type of focal epilepsy^1^. For most patients, seizures can be controlled with antiepileptic drugs (AEDs) and other treatments, but nearly a third of all cases remain resistant to therapy^2^. Drug resistant epilepsy is diagnosed when seizure freedom cannot be achieved after two trials of adequately chosen and tolerated AEDs^3^. The reasons why seizures in some patients respond well to AEDs while they do not in others remain poorly understood. Potential causes include genetic variants, drug target desensitization, and activity increase of drug efflux transporters ^4–7^. Associations between intractability, neurological network changes, and phenotype severity have been hypothesized^8–10^. Despite many advances, we lack comprehensive understanding of the effect of these alterations on the phenotype and subsequently, their contributions to treatment response remain largely unknown.

The lack of biomarkers to predict treatment response in focal epilepsy has long been noted^11^. While treatment-resistant TLE has been widely studied comparative studies of well-controlled and treatment-resistant patient populations are rare, yet informative. Recently, morphological changes of the corpus callosum in patients with early diagnosed focal epilepsy have been associated with treatment response^12^, and gray matter atrophy has been found to be more widespread in patients with pharmacoresistant mesial TLE than in well-controlled patients^13^. No differences of any functional imaging based biomarkers have been reported yet for intractability in epilepsy.

Resting state functional magnetic resonance imaging (Rs-fMRI) provides a powerful means to evaluate neural network connectivity and its alterations in epilepsy. Over the past decades, a number of Rs-fMRI studies of TLE have provided evidence for altered MRI-resting state functional connectivity (RsFC) patterns in TLE. Most studies found decreased RsFC between seizure network and ipsilateral structures of the temporal lobe, and increased RsFC between seizure network and contralateral temporal lobe structures^14–16^. Several studies, however, found decreased RsFC between seizure network and contralateral temporal lobe structures^17^ (reviewed in Centeno et.al, 2014^18^). None of these studies investigated differences between treatment-resistant and well-controlled patients. Hence, the effect of these network-alterations on the phenotype and their contributions to treatment response remain largely unknown.

Comparative imaging studies between treatment-resistant and well-controlled patients are difficult for a number of reasons. Epilepsy is a heterogeneous disorder and alterations in neural network organization that would relate to treatment response can easily be masked by functional connectivity alterations due to the anatomical diversity of seizure focus locations^19^. Moreover, Rs-fMRI is not routinely performed and hence, Rs-fMRI data, especially from patients with well-controlled epilepsy, are rare. Thus a large Rs-fMRI database needs to be screened to identify two patient sub-groups, treatment-resistant and well-controlled, that are otherwise comparable and share similar anatomical seizure focus localization.

Hypothesizing that intractable TLE patients exhibit larger differences to controls than tractable TLE patients, we reasoned that existing datasets, e.g. those of Voets et al^20^ comparing intractable TLE patients and healthy controls, could be utilized directly for power calculations and providing a reasonable estimate at necessary sample sizes. We considered a range of hypothetical tractable mean values (ranging from 10% and 90% of the intractable means reported in Voets et al^20^). Using these hypothetical tractable mean values in combination with the healthy control mean and SD values from Voets et al, we found group sizes of n=17 (for a 50% mean effect tractable/intractable) and n=6 (for a 90% mean effect tractable/intractable), respectively. Thus with a database large enough to contain such group sizes, it is possible to search for RsFC-based biomarkers.

We took advantage of an existing large imaging database at the NYU Langone Epilepsy Center, to study differences between tractable and intractable TLE patients, as well as control subjects. The database contains advanced imaging datasets from more than 600 patients and healthy controls, including detailed case characterizations, and Rs-fMRI data for a subset of individuals. Access to this large number of cases allowed us to focus on TLE cases and apply stringent criteria for the unbiased selection of patients with homogeneous conditions. To avoid potential confounds arising from differences in seizure focus lateralization, we focused on TLE patients only. Cases were separated into groups of right and left unilateral TLE patients and subsequently further divided into tractable and intractable cases. While this approach diminished the number of usable cases, we ultimately arrived at sufficiently large patient group sizes to test our hypothesis and show significant differences between patient groups.

We applied Rs-fMRI to explore a potential relationship between functional network changes and treatment response in TLE. We investigated differences in functional connectivity between regions of interest (ROI) within and beyond the epileptogenic network in patients with treatment-resistant and patients with well-controlled epilepsy. We targeted ROI-to-ROI connections of hippocampal and thalamic origin, as well as structures in proximity to the seizure network, including the amygdala, fusiform gyrus, and nucleus accumbens. We found robust and significant differences in functional connectivity patterns between TLE patients with controlled versus uncontrolled seizures. Thus, RsFC can potentially serve as a biomarker for treatment response in TLE.

## Methods

### Subjects

TLE patients and controls were recruited between 2009 and 2017 and consented to study protocols reviewed by the New York University’s Institutional Review Board. Patients were recruited via physician referral from the NYU Langone Epilepsy Center. Healthy controls were recruited through public advertisement. Imaging data from patients and healthy controls were selected by retrospective review of the NYU Langone Epilepsy Center’s MRI research registry, according to study criteria. Exclusion criteria comprised other neurological illnesses or serious medical problems (other than epilepsy in patients), estimated IQ within intellectual disability range (IQ < 70), current substance abuse, history of psychotic symptoms, and sensory problems. Healthy controls (HCs) were selected to match patients for age (+/- 5 years), gender and handedness. The absence of neurologic or psychiatric disease in HCs was assessed via semi-structured interviews. Patient data were only included when unilateral TLE had been diagnosed. Two independent, board-certified neurologists, both blinded to the study hypotheses, performed secondary review on an initial set of 34 eligible cases to confirm unilateral TLE, based on medical and neurologic evaluation, MRI, video-electroencephalography (EEG) monitoring and intracranial EEG, when available. Except for one well-controlled TLE patient, left lateralization was confirmed through MRI or EEG for all patients. Cases with unclear location of seizure focus, or bilateral or multifocal seizure activity were excluded from subsequent analyses. Remaining patient cases were divided into groups of right and left TLE. Patients with unilateral TLE were then further divided into two groups: one group consisting of cases of well-controlled TLE (cTLE) and a second group consisting of cases of treatment-resistant TLE (trTLE). Drug resistance was diagnosed by the attending neurologist and confirmed by the two independent reviewers, following Fisher et. al^3, 21^ criteria. Upon division of patient cases into cTLE and trTLE groups, only one right-localized cTLE case was found within our sample, and therefore, analyses were restricted to left-localized TLE cases. Division of right and left TLE cases was performed because the basis of treatment resistance is known to potentially vary between left and right lateralization^22^. Inclusion of uniformly left and unilateral TLE patients was performed to reduce noise inherent in inhomogeneous types of epilepsies and increases the methodological capability to detect treatment-response related differences between clinical groups. As alterations in neural network organization that would relate to treatment-response could easily be masked by anatomical diverse seizure focus locations, analyses were confined to unilateral and unifocal TLE cases. We included data from 28 individuals in the study: 13 HCs (19-54 years old, 4 female) and 15 TLE patients (19-63 years old, 10 female), with 7 cTLE cases and 8 trTLE cases, were included in subsequent analyses (For group information Table 1. For individual subject information see Supporting Table-S1. For detailed clinical information see Table-S3. A tree diagram outlining patient case screening and selection is provided in the SI section.

**Table 1:** Demographics and clinical characteristics

| **Table 1:** Demographics and clinical characteristics |  |  |  |
| --- | --- | --- | --- |
| **Variables** | **Controls**<br>**(N= 13)** | **trTLE**<br>**(N= 8)** | **cTLE**<br>**(N= 7)** |
| Gender (M/F) | 9/4 | 2/6 | 3/4 |
| Age (mean ± SD) | 36.2 ± 12.6 | 38.8 ± 6.4 | 29.1 ± 11.7 |
| Handedness (L/R) | 0/13 | 0/8 | 0/7 |
| Age of epilepsy onset (mean ± SD) | N/A | 17 ± 19.1 | 25.8 ± 14.1 |
| Duration of epilepsy (min – max, years) | N/A | 2 – 22* | 0 – 15* |
| Duration of epilepsy (mean ± SD, years) | N/A | 4.7 ± 7.8* | 3.3 ± 5.3* |
| Location and side of seizure focus (L/R) | N/A | 8/0 | 7/0 |
| Duration of follow up assessment (mean, years) | N/A | 4.5 | 3 |
| TLE = temporal lobe epilepsy, N/A = Not applicable for healthy controls, L/R = Left and right hemisphere. |  |  |  |
| *Duration of epilepsy unavailable for 4 patients, therefore ranges are reported in addition to the mean. |  |  |  |

### Image Acquisition

Participants were scanned on a 3T MRI Scanner (Siemens MAGNETOM, Allegra) using a single channel head coil. A T1-weighted anatomical image was collected (RT = 2.53s, TE = 3.25, flip angle = 7 degrees) in identical positioning to the following functional scan to allow for subsequent normalization. An echo planar resting state functional volume was acquired (AC-PC orientation, 64 × 64 matrix, 3 × 3 × 3 mm voxel size, 3mm slice thickness, echo planar imaging [EPI], 120 volumes, TR = 2s, TE = 25ms, flip angle = 90 degrees, 39 slices, field of view = 192mm^2^, duration = 540s). Participants were instructed to lie still in the scanner, keep their eyes open while focusing on a fixation cross on the screen, and not fall asleep.

### Preprocessing

Structural and functional raw scans were visually inspected for quality control and moved into the preprocessing pipeline. T1 scans were segmented into 164 regions of interest (ROIs) (Harvard-Oxford Atlas). Functional data preprocessing was carried out using the CONN toolbox^23^ (Version 17a.; http://www.nitrc.org/projects/conn). All volumes were realigned to the first functional scan, slice time corrected, co-registered to T1, segmented and normalized with the MNI-152 template. To avoid spurious results caused by increased sensitivity of resting state fMRI to head movement in high movement populations^24^ an aCompCor (anatomical component analysis correction) regression was applied and implemented using CONN toolbox. Compared to classical resting state preprocessing methods, aCompCor allows for robust data correction, additional movement and noise artifact reduction, while withstanding the need of global signal regression. This method delivers activation correlation results of high quality while allowing for analysis of negatively correlated ROI activation with a higher signal to noise ratio^25^.

In addition, resting-state time-series were linearly detrended, and bandpass filtered (0.008-0.09 Hz) to select for low frequency components and to reduce the confounding influence of respiratory (0.3 Hz) and cardiac (1 Hz) noise. Subsequently, data were spatially smoothed with a Gaussian kernel (FWHM 6mm). Functional scans were additionally quality controlled: functional connectivity was controlled for skewing, high movement volumes (above 1 mm) were scrubbed and the mean image intensity and standard deviation were reviewed for outliers, using an artifact detection tool (ART, build into CONN v17a.). Volumes containing measurements above the set thresholds were excluded from further analysis (between 2 and 13 volumes for 7 controls, 2-17 volumes for 13 trTLE, and 2-13 volumes for 4 cTLE).

To control for spurious results due to group differences in movement we compared the mean inter-volume translation of healthy controls (0.034 ± 0.015 SD), treatment-resistant trTLE (0.039 ± 0.016 SD), and well-controlled cTLE (0.037 ± 0.029 SD) and found no significant difference. Furthermore, all scans were manually quality controlled by visual inspection before and after preprocessing to ensure good scan quality and to avoid inclusion of artifact containing data.

The whole brain was parcellated into 164 structurally homogenous regions of interest (ROIs), in accordance with the FSL Harvard-Oxford atlas for the grey matter as well as subcortical regions. For the default mode network 4 structurally homogeneous ROIs were taken from the literature^26^. ROIs were derived from the parcellated structural T1 scan of each subject and normalized within the Montreal Neurological Institute 152 Space (MNI152 space). Finally, connectivity data were computed using SPM and CONN software packages and the adjacency matrix, reflecting ROI-to-ROI activation, and signal strength correlation, was calculated and output values were used for analyses (Figure 1).

**Figure 1.**
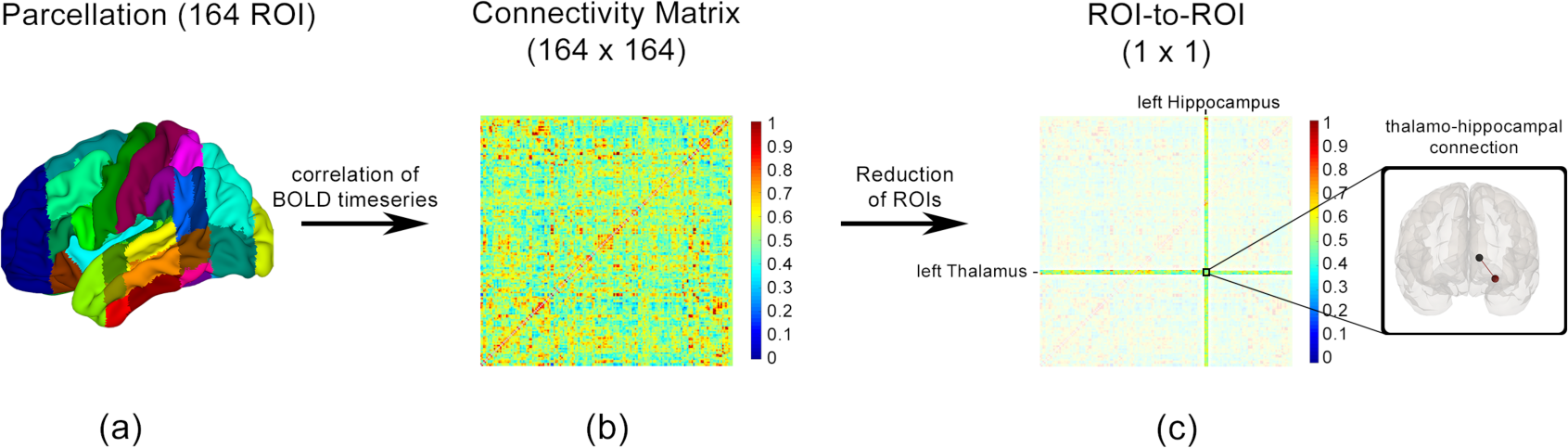
Processing steps per case, a) parcellation of T1-weigthed anatomical MRI data into 164 pre-defined (Harvard Oxford Atlas) regions of interest (ROIs), b) mean Fisher-transformed Z-scores of Resting State Functional Connectivity (RsFC) correlation between all 164×164 ROI-to-ROI connections shown in one connectivity matrix, c) extraction of RsFC connectivity Z-scores of interest.

### Region of interest (ROI) selection

We investigated ROI-to-ROI connections between separate brain regions. Of all possible connections (164×164), we restricted our analyses to a specific set of 35 connections (see SI for detailed table). Past literature of trTLE suggested predictability of seizure focus lateralization through measurement of RsFC between the thalamus, hippocampus and amygdala^27^. We therefore selected thalamic connections as well as temporal regions (hippocampus, amygdala, fusiform cortex) that were expected to be in proximity to the assumed seizure focus. In addition, RsFC between ROIs from well-studied networks (default mode, salience and attention network) as well as ROIs unlikely to be altered by TLE, such as cerebellum and brainstem, were included for control measures.

Using these 35 connections, we investigated individual differences as well as differences between the three groups (Supporting Table-S2).

Lastly, we separately tested an additional set of seven ROI-to-ROI connections which have previously been shown to be predictive of postsurgical failure^28^. These six connections include: left inferior temporal - to - left pallidum, left lateral occipital - to - left amygdala, left anterior middle temporal gyrus - to - left putamen, left superior temporal region - to - left anterior insula, left superior temporal - to - left anterior insula, and left anterior cingulate - to - left hippocampus, and left superior frontal gyrus – to – left medial temporal lobe (as an approximation of the uncinate fasciculus connections) (Supporting Table-S2b).

### Statistical analysis

Analyses were performed using IBM SPSS v22. ANCOVA was applied to detect differences in RsFC between the three groups with age at time of scanning as a covariate. Post-hoc Tukey t-tests with Bonferroni multiple comparison correction were used on significant omnibus group results. In addition, to correct for multiple hypothesis testing of 35 ROI-ROIs, results significantly differing between trTLE and cTLE groups were corrected once more for multiple comparison (Bonferroni).

To probe the combined measurements’ power to distinguish between cTLE and trTLE, we modeled significant thalamo-hippocampal connections in a mixed-effects model for repeated measures. In the model, group and hemisphere were included as fixed effects and subject was included as a random effect to account for the spatial repeated measurements by hemisphere. For this analysis we utilized the R software tool and confirmed the results by using SAS software suite.

To visualize the relationship between the significant RsFC values within our sample, we plotted the connectivity measures for both thalamo-hippocampal connections (left and right) in a two-dimensional canonical space. These individual results can be clustered according to our clinical groups. By computing the distance between these cluster-clouds (which we will refer to as the Mahalanobis distance (MD)), we arrive at a unitless measurement of difference between the groups. This unit-independent, scale-invariant, and correlation independent approach allows for the measurement of distance between a point (defined by the center of the cluster of one group) and a distribution (defined by the distribution of the other group) in multidimensional space. We chose to use this measure because of its reliability when comparing two or more dependent variables as in the case of bilateral functional connectivity between two brain regions. If the two groups resemble each other in their bilateral thalamo-hippocampal RsFC, then we expect a small MD between the centers of the cluster clouds. If on the other hand the two groups can be distinguished along the measure of those connections, then we expect a larger MD.

Additionally, to account for our small sample size we applied randomization and bootstrapping methods to the MD approach. We used the true, non-randomized, clinical groups and bootstrapped the MD by running 10,000 iterations (sampled with replacement). To judge reliability of the MD results (original and bootstrapped), we performed a second bootstrapping using randomized groups. We randomly assigned each subject to one of two arbitrary groups and then calculated the MD between these groups (again 10,000 runs, sampled with replacement). By bootstrapping true MD results and comparing these with results from randomized groups we are able to increase the reliability of our results within the statistical limitations of our sample size.

## Results

We investigated RsFC across 35 ROI-to-ROI connections of hippocampal and thalamic origin, as well as amygdala, fusiform gyrus, nucleus accumbens, default mode, salience, and attention network, brainstem and cerebellar ROIs (for a full list of RsFC results see Supporting Table-S2). Of the 35 connections investigated, only two significantly differed between trTLE and cTLE: thalamo-hippocampal RsFC in both hemispheres, while positively correlated in both cTLE patients and HCs, was negatively correlated in trTLE patients (Figure 2). On a group level, thalamo-hippocampal RsFC could distinguish cTLE from trTLE. Functional connectivity between left thalamus and left hippocampus differed between the two patient groups (ANCOVA, *F* (2,24) = 13.2, p < 0.001, r = 0.51) (Figure 2a). Post-hoc Bonferroni correction revealed that left thalamo-hippocampal RsFC was decreased in patients with trTLE when compared to patients with cTLE (p < 0.01, for ANCOVA). This result remained significant after correcting for multiple hypotheses testing (p=0.005). Left thalamo-hippocampal RsFC in HCs did not differ from cTLE but was distinct from trTLE patients (p < 0.01, Bonferroni corrected). A similar, but statistically weaker pattern was found when RsFC between right thalamus and right hippocampus was investigated. Functional connectivity between those regions was decreased in trTLE patients (ANCOVA, *F* (2,24) = 3.67, p = 0.04, r = 0.26) (Figure 2b). However, post-hoc Bonferroni correction (p=0.063, for ANCOVA) revealed that right thalamo-hippocampal RsFC was not significantly decreased in people with trTLE when compared to people with cTLE. These results demonstrate differences in thalamo-hippocampal RsFC between people with well-controlled and treatment-resistant left TLE that were more pronounced in the hemisphere ipsilateral to the seizure onset.

**Figure 2.**
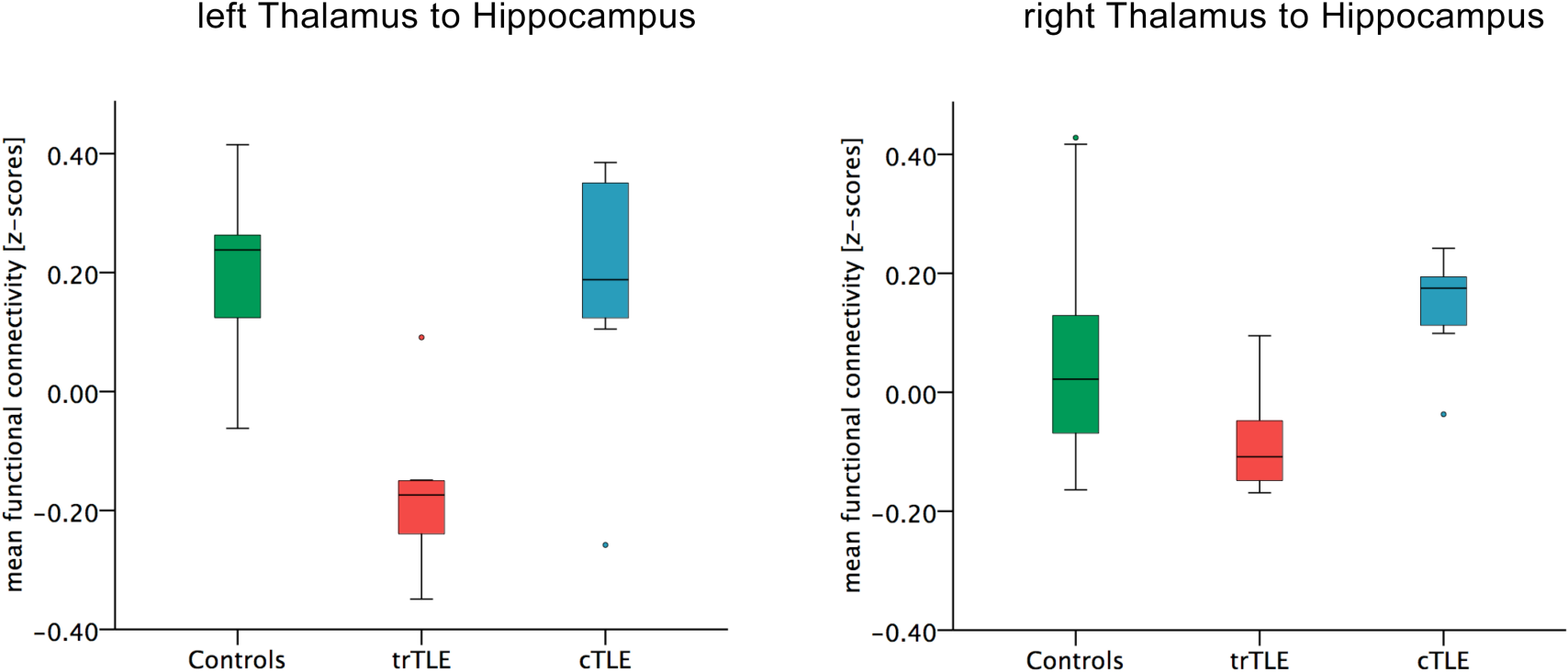
Mean Fisher-transformed Z-scores of Resting State Functional Connectivity (RsFC) correlations between the left thalamus and left hippocampus, and between the right thalamus and right hippocampus in healthy controls (Controls, green, n=13), treatment-resistant left TLE patients (trTLE, red, n=8), and well-controlled left TLE patients (cTLE, blue, n=7). Error bars indicate the 95% confidence interval.

Following up on the differences in thalamo-hippocampal RsFC between trTLE and cTLE groups, we were interested in the consistency of these effects from subject to subject. Figure 3 shows individual left thalamo-hippocampal RsFC measurements. Strength and directionality of this connection varied across subjects but the general qualitative pattern of group differences was apparent. All but one HC, and all but one cTLE case, exhibited positive functional connectivity, while all but one trTLE patient exhibited negative connectivity. Functional thalamo-hippocampal connectivity patterns in cTLE patients overlapped with those in HCs. These results underscore that, despite individual differences in the strength of left thalamo-hippocampal RsFC, the directionality (i.e., positive versus negative) reliably differentiated patients with cTLE (and HCs) from patients with trTLE at the individual level.

**Figure 3.**
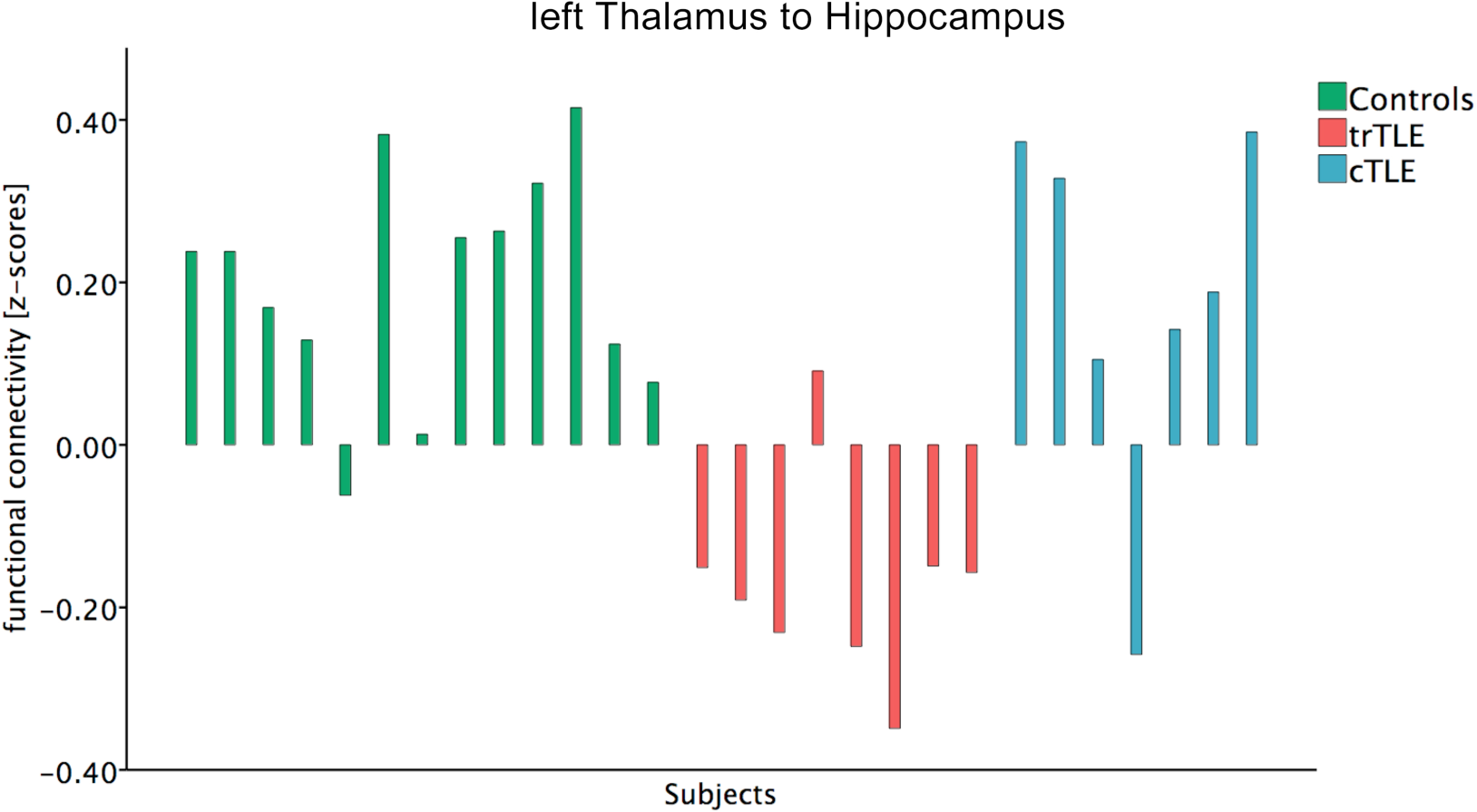
Subject level analysis of functional connectivity shows individual differences in Fisher-transformed Z-scores of Resting State Functional Connectivity (RsFC) correlations between the left thalamus and the left hippocampus. Healthy controls (Controls, green, n=13), treatment-resistant left TLE patients (trTLE, red, n=8), well-controlled left TLE patients (cTLE, blue, n=7).

Group-level and individual-subject level analyses (Figures 2 and 3) suggest that thalamo-hippocampal RsFC might serve as a biomarker for trTLE. To explore this possibility further, we plotted right and left thalamo-hippocampal RsFC for each patient in a scatter plot (Fig. 4). We characterized cTLE and trTLE group cluster separation with the Mahalanobis distance and assessed its significance using bootstrapping (see Methods). The distance between the true groups was significantly higher (21.1 ±18.7) than the distance between the randomized groups (0.7 ±1.0) (Figure 5). Thus, left and right thalamo-hippocampal RsFC measures allowed for a differentiation between cTLE and trTLE. Moreover, we combined measurements of right and left hemispheric thalamo-hippocampal RsFC to test for overall difference between the two groups. This mixed effects model found the group effect to be significant at p=0.0004, while there was no significant group-by-hemisphere interaction (p=0.14). Together, these results show a highly significant difference in thalamo-hippocampal RsFC between the two patient groups.

**Figure 4.**
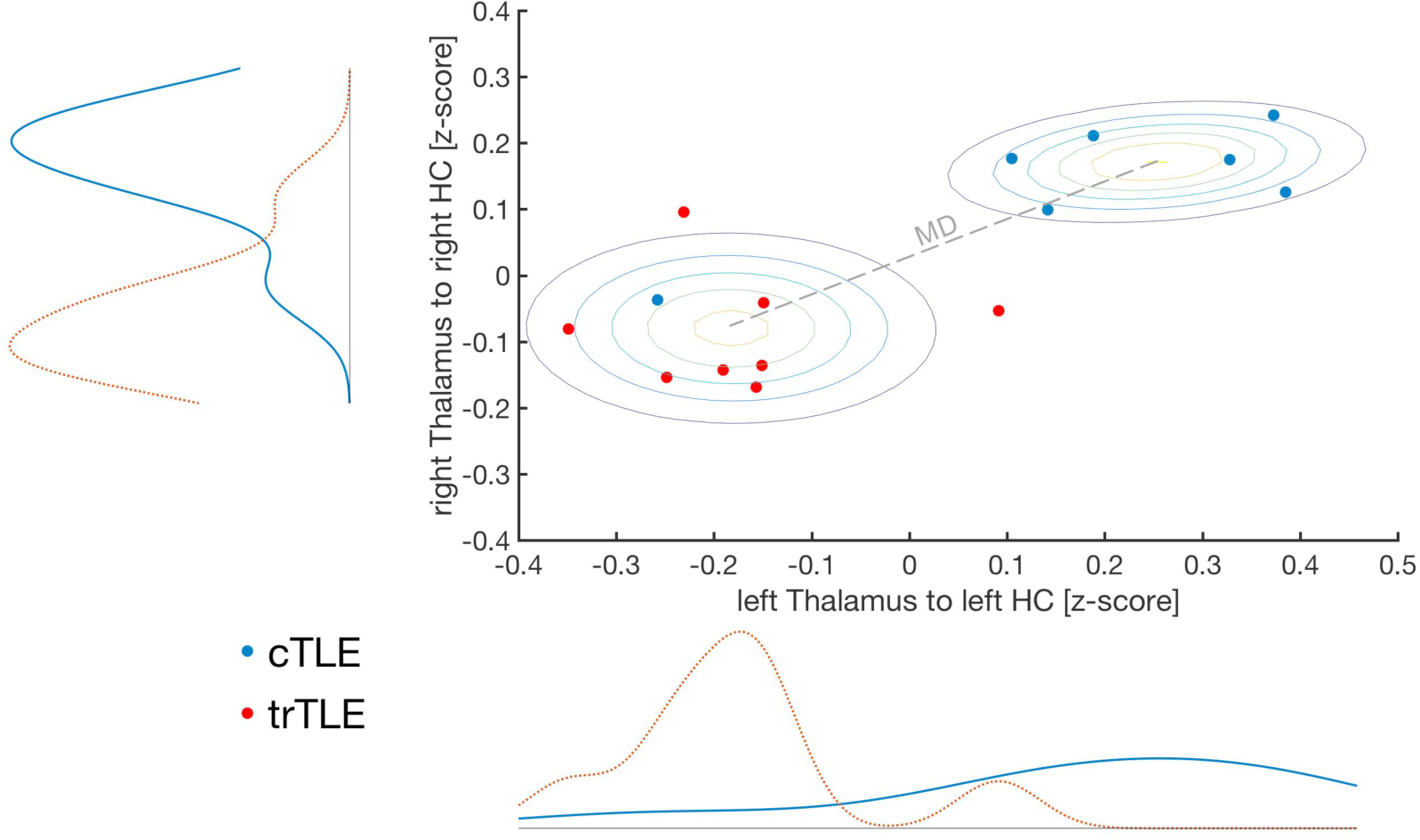
Scatterplot showing individual measurements in 2-dimensional feature space, containing of ipsilateral thalamo-hippocampal RsFC measures bilaterally (left Thalamus to left Hippocampus (HC)). Showing two clearly distinguishable clusters of well-controlled and treatment-resistant left-TLE cases, with one clear outlier, one medically tractable case falling among the cluster of treatment-resistant cases. The Mahalanobis distance (MD) is computed between the two cluster-centers, seen here visualized with a Gaussian mixture distribution. Contour lines are added representing the probability distribution function (min-max).

**Figure 5.**
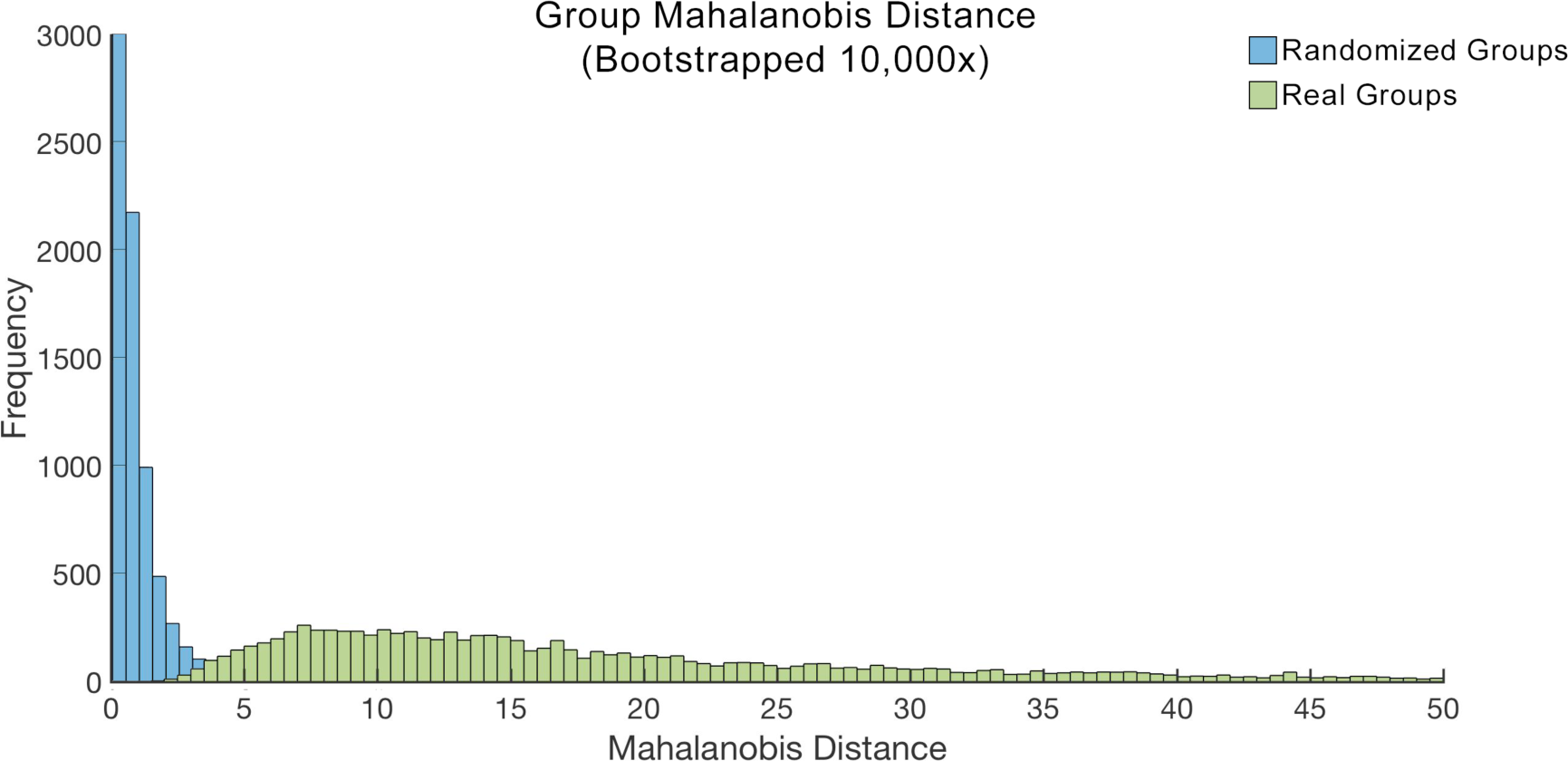
The Mahalanobis distance between the two group clusters (for bilateral thalamo-hippocampal connectivity, see Figure 4) was randomized (10,000x bootstrap – with replacement). Randomization was performed once on subject data assigned to the correct groups (trTLE and cTLE, green) and once when subject data was randomly assigned to one of the two groups (blue). The histogram results of the bootstrapping are shown for both real and random groups. With random groups approaching zero, or no significant Mahalanobis distance. Real group distances ranged from 1 to over a 100 (shown here for the range 0 to 50).

Separately, we analyzed seven functional connections that have previously been shown to predict post-surgical outcome (see Methods). Of those seven, none were significantly different between groups using the same ANCOVA method as described above (duration of epilepsy as covariate, multiple comparison correction).

## Discussion

Using functional connectivity correlates of treatment response in TLE, we discovered that thalamo-hippocampal RsFC was abnormal in strength and direction in trTLE relative to cTLE and HCs. The differences were large and robust to multiple comparison correction, as well as consistent across different analytic methods. These results implicate thalamo-hippocampal RsFC measures as a potential biomarker for TLE treatment response.

This is the first time RsFC differences between well-controlled and treatment-resistant TLE patients have been found. Previous studies investigated RsFC patterns in trTLE patient populations alone, while comparison of RsFC across different brain regions in trTLE versus cTLE has never been studied. We found a difference between left trTLE and left cTLE in RsFC strength and directionality that was localized to thalamo-hippocampal pathways and was more pronounced in the left hemisphere and thus, ipsilateral to the seizure focus. Thus, RsFC abnormalities in trTLE appear to be spatially localized rather than global or particularly pronounced on the side ipsilateral to seizure onset.

Surprisingly, we found reduced, not enhanced, thalamo-hippocampal connectivity as a signature of trTLE. The mechanisms driving reduced thalamo-hippocampal connectivity are unclear but may involve thalamic nuclei such as the medial dorsal thalamic nuclei and the nucleus reticularis. The medial dorsal thalamic nuclei are critical for limbic seizure propagation^29^, and unilateral stimulation of this nucleus in kindling models suppresses limbic seizures^29, 30^. The nucleus reticularis forms a shell of inhibitory cells around the thalamus; over-activation of these inhibitory neurons in trTLE could reduce functional connectivity with the hippocampus. Mechanistic work in animal models should determine the plausibility of such a mechanism for explaining long-range functional connectivity abnormalities.

Our methods do not allow us to distinguish cause from consequence. Altered thalamo-hippocampal connectivity could result from persistent seizure activity or an etiological factor that contributes to treatment-resistant seizures. Frequent seizure activity could change network organization. This hypothesis is supported by work showing modulating effects of seizure activity on functional connectivity both within and beyond seizure networks^16^, and increasing effects with time of chronic seizure exposure^31^. Alternatively, aberrant thalamo-hippocampal RsFC may contribute to intractability in trTLE. For example, altered connectivity might contribute to the difference in seizure frequency that occurs even before initiation of therapy in patients who subsequently show inadequate response to treatment^2^. If this is the case, then altered functional connectivity might impact the success of drug treatment trials. To test these hypotheses, future efforts should be focused on routine acquisition of RsFC fMRI data from newly diagnosed and treatment naïve patients for longitudinal studies.

The finding of a specific alteration in trTLE connectivity between two brain regions has sufficiently important implications to warrant future studies. Longitudinal observation of changes in thalamo-hippocampal RsFC from the time of seizure onset through the course of pharmacological treatment can elucidate the directionality of the relationship between RsFC characteristics and treatment response. Should thalamo-hippocampal RsFC abnormalities prove to be the consequence of persistent seizures, then future work should determine the degree to which this contributes to cognitive and psychiatric comorbidities in TLE. Thalamo-hippocampal RsFC could be an early predictor of treatment response, making this a valuable means to identify patients who would benefit from adjusted treatment regimen or earlier surgical consideration.

Our findings provide the first evidence for differences in neural network connectivity between well-controlled and treatment-resistant TLE patients. The functional network alterations, whether as cause or consequence of trTLE, show promise as a putative biomarker. These results suggest future multi-center studies. Future studies with longer resting-state scans can improve signal to noise ratios and potentially increase diagnostic power and inclusion of accompanying structural connectivity measures such as diffusion weighted imaging based tract tracing would be valuable for modeling the structural correlates of RsFC abnormalities. Changes in neural fiber structure density between thalamus and hippocampus in TLE^32, 33^, and measures of functional connectivity and fiber density can predict seizure focus lateralization in TLE^27, 32^.

Moreover, cytoarchitectonical abnormalities have be visualized in TLE patients under utilization of diffusional kurtosis imaging (DKI)^34^, and structural brain connectome measures both, pre- and postoperatively are associated with postsurgical seizure freedom in TLE^28, 35^. Importantly, analyses of RsFC between seven ROI-to-ROI connections that had previously been implicated to be predictive of surgical failure revealed no differences between groups. These results indicate that pharmacological treatment resistance and failure of surgical control are two separate topics that are not necessarily related. Potential differences within the trTLE group could not be investigated. Due to the scarce number of cases and incomplete information on post surgical outcome within the trTLE group comparative analyses and correlation of functional connectivity measures with post surgical outcome could not be performed. Nonetheless, future studies should examine whether differences in functional connectivity between treatment resistant and well-controlled TLE patients are associated with altered structural connections.

In sum, our findings of thalamo-hippocampal RsFC alterations in trTLE patients could lead to the identification of new pathways to treatment resistance, guide drug target and biomarker discovery, which could ultimately result in improved diagnosis and treatment for people with TLE.

## Acknowledgements

We would like to thank Dr. Yifat Prut and Dr. Heath Pardoe, for helpful discussions, valuable input and comments on the project. We would also like to thank Caroline Jiang as well as Dr. Roger Vaughan for their valuable feedback on the statistics sections.

## Conflicts of Interest and Affirmation

The authors report no conflicts of interest.

## Author Contributions

Study concepts/study design or data acquisition or data analysis/interpretation, all authors; manuscript drafting or manuscript revision, all authors; approval of final version of submitted manuscript, all authors; literature research C.P., P.B., K.B, P.D.; clinical case review, M.H. P.D.; statistical analysis, C.P., P.B., S.S.; and manuscript editing, all authors.

